# Astrocytic uptake of posttranslationally modified amyloid-β leads to endolysosomal system disruption and induction of pro-inflammatory signaling

**DOI:** 10.1101/2023.09.04.556200

**Authors:** S Wirth, A Schlößer, A Beiersdorfer, M Schweizer, C Lohr, KM Grochowska

**Author notes:** To whom all correspondence should be sent at: Leibniz Group ‘Dendritic Organelles and Synaptic Function’, Center for Molecular Neurobiology, ZMNH, University Medical Center Hamburg-Eppendorf, 20251 Hamburg, Germany. These authors have contributed equally (S Wirth, A Schlößer).

## Abstract

The disruption of astrocytic catabolic processes contributes to the impairment of amyloid-β (Aβ) clearance, neuroinflammatory signaling, and the loss of synaptic contacts in late-onset Alzheimer’s disease (AD). While it is known that the posttranslational modifications of Aβ have significant implications on biophysical properties of the peptides, their consequences for clearance impairment are not well understood. It was previously shown that N-terminally pyroglutamylated Aβ3(pE)-42, a significant constituent of amyloid plaques, is efficiently taken up by astrocytes, leading to the release of pro-inflammatory cytokine tumor necrosis factor α (TNFα) and synapse loss. Here we report that Aβ3(pE)-42, but not Aβ1-42, gradually accumulates within the astrocytic endolysosomal system, disrupting this catabolic pathway and inducing formation of heteromorphous vacuoles. This accumulation alters lysosomal kinetics and lysosome-dependent calcium signaling, and upregulates lysosomal stress response. These changes correlate with the upregulation of glial fibrillary acidic protein (GFAP) and increased activity of nuclear factor kappa-light-chain-enhancer of activated B cells (NFᴋB). Treatment with a lysosomal protease inhibitor, E64, rescues GFAP upregulation, NFᴋB activation, and synapse loss, indicating that abnormal lysosomal protease activity is upstream of pro-inflammatory signaling and related synapse loss. Collectively, our data suggest that Aβ3(pE)-42-induced disruption of the astrocytic endolysosomal system leads to cytoplasmic leakage of lysosomal proteases, promoting pro-inflammatory signaling and synapse loss, hallmarks of AD-pathology.

## Introduction

The processes underlying the onset and progression of Alzheimer’s disease (AD) are substantially driven by glial signaling (Haass & Selkoe, 2022; Knopman et al, 2021; Selkoe & Hardy, 2016). Recent scientific evidence has shed light on the role of microglia in the disease, while the role of astrocytes remains less understood (De Strooper & Karran, 2016; Patani *et al*, 2023). Astrocytes can contribute to disease onset and progression by modulating neuroinflammatory signaling and debris clearance (De Strooper & Karran, 2016; Haass & Selkoe, 2022; Heneka *et al*, 2015; Knopman *et al*., 2021; Patani *et al*., 2023; Rodríguez-Arellano *et al*, 2016). The latter is especially important in the context of abnormal amyloid-β (Aβ) accumulation, a hallmark of AD (De Strooper & Karran, 2016; Haass & Selkoe, 2022; Knopman *et al*., 2021; Selkoe & Hardy, 2016). According to the prevalent hypothesis, defective Aβ clearance results in enhanced concentration of soluble oligomeric Aβ species, which in turn is the key factor leading to AD-related synaptic loss and neurotoxicity (De Strooper & Karran, 2016; Haass & Selkoe, 2022; Selkoe & Hardy, 2016). Diverse Aβ peptides have been described that arise from different processing of amyloid precursor protein (APP) and numerous posttranslational modifications (Kummer & Heneka, 2014; Polanco *et al*, 2018; Selkoe & Hardy, 2016). Among them N-terminally truncated, pyroglutamylated Aβ, Aβ3(pE)-42, is prominently present in post-mortem human brain (Harigaya *et al*, 2000; Jawhar *et al*, 2011; Saido *et al*, 1995). Compared to Aβ1-42, Aβ3(pE)-42 displays a higher stability and aggregation propensity (D’Arrigo *et al*, 2009; Jawhar *et al*., 2011; Nussbaum *et al*, 2012; Saido *et al*., 1995; Schlenzig *et al*, 2009). In addition, pyroglutamate formation enhances Aβ3(pE)-42 hydrophobicity and interaction with lipid membranes (Gillman *et al*, 2014; Harigaya *et al*., 2000; Lee *et al*, 2014; Portelius *et al*, 2010). In previous work we revealed that contrary to Aβ1-42, Aβ3(pE)-42 is taken up and accumulates in astroglia but not neurons, which in turn induces the release of pro-inflammatory tumor necrosis factor α (TNFα) that drives synaptic loss (Grochowska *et al*, 2017). However, the link between astrocytic clearance of Aβ3(pE)-42 and induction of pro-inflammatory, synaptotoxic signaling is at present unclear.

In this study, we decipher the cellular mechanisms that underlie the formation of Aβ3(pE)-42 accumulation in astrocytes. We demonstrate that Aβ3(pE)-42 oligomers gradually accumulate, induce disruption of the endolysosomal system and formation of larger single-membrane vacuolar structures, indicating profound impairment in astrocytic catabolic mechanisms. Furthermore, we show that Aβ3(pE)-42 accumulation promotes impairment in lysosomal Ca^2+^ signaling, induces a lysosomal stress response, and, as a consequence, upregulation of glial fibrillary acidic protein (GFAP) and increased activity of the nuclear factor kappa-light-chain-enhancer of activated B cells (NFᴋB). Collectively, our findings suggest that Aβ3(pE)-42-driven disruption of the astrocytic endolysosomal system underlies the onset of pro-inflammatory signaling leading to synapse loss.

## Methods

### Animals

All animal experiments were carried out following the European Communities Council Directive (2010/63/EU) and approved by the local authorities of the city-state of Hamburg (Behörde für Gesundheit und Verbraucherschutz, Fachbereich Veterinärwesen) and the animal care committee of the University Medical Center Hamburg-Eppendorf. For primary, astrocytic culture P0 JAX C57BL/6 mice or B6;129-Gt(ROSA)26Sortm1(CAG-cas9*,-EGFP)Fezh/J crossed with Cre-driver line (Schwenk *et al*, 1995) mice were used. For immunohistochemical studies, brain cryosections of 24 weeks old TBA2.1 mice were used (Alexandru et al., 2011). TBA2.1 mice were a kind gift from Probiodrug, now Vivoryon Therapeutics N.V. Wistar rat embryos (E18) of mixed sex were used for primary neuronal cultures.

### Primary murine astrocytic culture

For the preparation of primary glial cells, the cortex of P0 mice was fragmented in Hank’s Balanced Salt Solution (HBSS) at room temperature (RT) and then incubated for 15 min with 0.25% trypsin at 37°C. Subsequently, the tissue was fragmented in the presence of Deoxyribonuclease I (DNase I) (Roche, Cat.: #11284934001). The cells were grown for 10-14 days in the Dulbecco’s Modified Eagle’s Medium (DMEM) supplemented with 10% fetal bovine serum (FBS), 1% L-glutamine (Thermo Fisher Scientific, Cat.: #25030149). DIV10-14 astrocytes were split and used for experiments. For immunocytochemistry staining, coverslips coated with poly-L-lysine (PLL) (100 mg/l in 100 mM boric acid pH 8.4) were used. PLL-coated aclar film (Ted Pella, Inc., Cat.: # 10501-10) was used for the transmission electron microscopy (TEM) experiment. The preparation is devoid of microglia.

### Primary hippocampal cultures

Primary hippocampal cultures were prepared according to a previously published protocol (Grochowska *et al*, 2023). Briefly, hippocampi were dissected from Wistar rat embryos (E18) and then washed five times in ice-cold HBSS. The tissue was incubated at 37°C in 0.25% trypsin for 15 min. Dissociated cells were plated at a density of 60 000 cells per well in DMEM medium supplemented with 10% fetal calf serum (FCS) and 2 mM glutamine in 12-well dishes on PLL-coated glass coverslips. Cultures were maintained at 37°C, 5% CO_2_, and 95% humidity. The cultures contain astroglia and neurons but no microglia.

### Preparation and characterization of amyloid oligomers

Human Aβ1-42 (Anaspec, Cat.: #AS-20276), Aβ3(pE)-42 (Anaspec, Cat.: #AS-29907), and FAM-Aβ3(pE)-42 (Eurogentec, custom-made) were prepared as previously described (Grochowska *et al*., 2017; Klein, 2002). Briefly, for Aβ1-42 oligomeric preparation, the peptide film was dissolved in 2 μl dimethyl sulfoxide (DMSO) and sonicated for 5 min. Afterwards, the F12 medium was added to the final concentration of 50 μM and the peptides were sonicated for 10 min. The peptides oligomerized for 24 h at 4°C and were centrifuged for 10 min and 14000 rcf at 4°C prior to treatment. Human Aβ(pE)-42 and FAM-Aβ3(pE)-42 peptides were oligomerized according to the published protocol (Nussbaum et al., 2012). The peptides were dissolved in 0.1 M NaOH, and diluted in neurobasal medium (Gibco, Cat.: #21103049) to the final concentration of 5 μM. 1 M HCl was used to adjust the pH. Subsequently, the peptides oligomerized for 24 h at 37°C. A final concentration of 500 nM was used for Aβ3(pE)-42 oligomer treatment. The oligomerization was tested by Western blot (WB). All oligos were used at a final concentration of 500 nM, the treatment duration is indicated in the figure legends.

### Cell transfection and treatment

All drugs and constructs used in the study can be found in the Table S1. The primary astrocytes were transfected one day before treatment with Lipofectamine 2000 (ThermoFisher Scientific, Cat.: #11668-019), according to the manufacturer’s instructions. The final concentrations of drugs used in the study: 10 μM of E-64 (Sigma-Aldrich, Cat.: #E3132-1MG), 60 nM of *t*-NED19 (Bio-techne, Cat.: #3954). The duration of treatments is indicated in the figure legends.

### Immunocytochemistry

All antibodies used in this study can be found in the Table S1. The coverslips were fixed for 10 min in 4% paraformaldehyde (PFA)/4% sucrose at RT. The cells were then washed three times for 5 min with phosphate buffered saline (PBS) and, permeabilized with 0.2% Triton X-100 in PBS for 10 min at RT, and subsequently incubated for 1 h at RT in blocking buffer (2% glycine, 2% bovine serum albumin (BSA), 0.2% gelatin, and 50 mM NH_4_Cl), followed by overnight incubation at 4°C with primary antibodies diluted in blocking buffer. Next, the samples were washed and incubated with the secondary antibody dilution for 1 h at RT. Finally, the sample was washed and the coverslips mounted with Mowiol 4-88. The coverslips were stored in the dark at 4°C.

### Immunohistochemistry

All antibodies used in this study can be found in the Table S1. Coronal cryosections from 24-week-old TBA2.1 mice and wild-type (WT) littermates were used for immunohistochemical staining. For antigen retrieval, free-floating sections were incubated with 10 mM sodium citrate (pH=9) for 30 min at 80°C. Samples were permeabilized with 0.3% Triton X-100 for 10 min at RT. Next, sections were incubated in blocking solution at RT for 30 min. Then, the primary antibody diluted in blocking solution was added and incubated at 4°C for 72 h. Subsequently, the sections were washed three times with PBS at RT for at least 1 h. The secondary antibody was prepared at a concentration of 1:500 in blocking solution and incubated for 2 h at RT. The sections were washed three times again with PBS. The fluorescent dye 4’,6-Diamidino-2-phenylindole (DAPI) was diluted in PBS (1:400) for nuclear staining. After 30 min of incubation in DAPI solution, sections were briefly washed with PBS followed by distilled water. Sections were mounted on a coverslip using Mowiol 4-88 and stored in the dark at 4°C.

### Confocal microscopy and data analysis

Samples were imaged with a Leica TCS SP5 (HCX PL APO 63 ×/1.40 NA objective) or Olympus BX61WI confocal microscope (Nikon P-Apo 60×/ 1.40 NA objective) controlled by software provided by manufacturers (Leica LAS AF software and Olympus FV10-ASW respectively). Selected region was scanned with 405, 488, 568 and 635 nm laser lanes (12-bits, 80 × 80 nm pixel size, 700 Hz, Z-step 0.42 – 1 µm).

For the experiments comparing mean staining intensities between treatment, the imaging parameters were set according to the control group and kept consistent for all scanned groups. Fiji/ImageJ (Schindelin *et al*, 2012) was used for image analysis.

To measure the GFAP or NF-кB reporter fluorescence intensities, the average mean intensity of three z-stacks (depth 0.84 μm) was used and the intensities within a cell (defined by GFAP signal) were measured. For the measurement of association of the Aβ3(pE)-42 with markers of the endolysosomal compartment, the average mean intensity of five z-stacks (depth 1.68 μm) was used and regions of interest (ROIs) were segmented using thresholding functions.

For synaptic density measurements, the maximum projection of three z-stacks was created. The synapses were defined as the overlap or apposition signal from the postsynaptic density protein 95 (PSD95) and synaptophysin 1. The analysis was done on the primary, proximal dendritic segment. The number of synapses was normalized to the length of analyzed dendritic stretch.

Stained cryosections were acquired with a pixel size of 1 μm. A maximum projection of ten z-stacks was used to measure the mean intensity of GFAP and ionized calcium-binding adapter molecule 1 (Iba1). All mean intensities were normalized to the averaged value of WT section.

### Cell viability assay

Astrocytes were seeded in the flat-bottom, white, 96-well plates. A luciferase-based RealTime-Glo MT Cell Viability Assay (Promega, Cat.: #G9711) was performed according to the manufacturer instructions. The luminescence was measured using Tecan Spark 10M (Tecan).

### Lysosomal mobility assay

Live imaging of primary astrocytes was performed with a Nikon Eclipse Ti-E controlled by VisiView software (Visitron Systems), with 100× objective (Nikon, CFI Plan Apochromat Lambda 100×/ 1.45) equipped with an integrated Nikon Perfect Focus system. TIRF images were acquired with a spinning TIRF system (iLAS2, Gattaca Systems). The sample was imaged in Tyrode’s buffer in the imaging chamber and imaged at an acquisition frequency of 1 Hz for 3 min at 37°C and 5% CO_2_. The samples were incubated with 0.5 µM SiR-lysosome (Spirochrome, Cat.: #SC012) for 60-120 min prior to imaging. For analysis the Velocity Measurement Tool of Fiji/ImageJ (Schindelin *et al*., 2012) was used.

### Astrocytic Ca^2^+ imaging

Confocal Ca^2+^ imaging of cultured astrocytes was performed according to (Stavermann *et al*, 2012). Primary astrocyte cultures were loaded with the 5 µM chemical calcium indicator Rhod-2 AM (Abcam, Cat.: #ab142780) in standard artificial cerebrospinal fluid (ACSF; in mM: 120 NaCl, 2.5 KCl, 1 NaH_2_PO_4_x2H_2_O, 26 NaHCO_3_, 2.8 D-(+)-glucose, 1 MgCl_2_, 2 CaCl_2_). Cultures grown on PLL-coated glass coverslips were placed in the experimental bath and fixed using a U-shaped platinum wire. The samples were continuously perfused with ACSF and gassed with carbogen (95% O_2_, 5% CO_2_) to maintain the pH of 7.4 and to supply oxygen. All drugs were applied via the perfusion system. The agonist adenosindiphosphate (ADP, AppliChem #A0948, 25 µM in ACSF) was applied for 30 s. The antagonist *t*-NED19, 60 nM (bio-techne, #3954) was applied for 10 min prior to agonist application. Changes in cytosolic calcium concentration were detected by the fluorescence of Rhod-2 (excitation: 543 nm) using a confocal microscope (eC1, Nikon). Images were acquired at a time rate of one frame every 3 s. To analyze changes in cytosolic Ca^2+^, ROIs were defined using Nikon EZ-C1 software. Changes in Ca^2+^ were analyzed throughout the experiment as relative changes in Rhod-2 fluorescence (ΔF) in respect to the base line fluorescence, which was normalized to 100%. All values are stated as mean values of fluorescence amplitude.

### Immunoelectron microscopy

For ultrastructural analysis primary astrocytes were treated with Aβ3(pE)-42 oligomers for 24 h. They were fixed with a double strength fixative (4% PFA and 2% Glutaraldehyde (GA) mixed with medium 1:1) at RT for 15 min. Thereafter the cells were fixed with 4% PFA and 1% GA only, overnight.

Pre-embedding immunolabelling was performed as following: Dissociated astrocytes were fixed with PB (phosphate buffer) containing 4% PFA and 0,1% glutaraldehyde. 2.3 M sucrose was used for cryoprotection. Penetration of immunoreagents inside the cells was obtained by two freeze-thaw cycles in liquid nitrogen. After washing in PBS, 10% horse serum (PS) containing 0.2% BSA was used to block unspecific binding sites for 15 min. Primary antibodies (for details see Table S1) were applied overnight in PBS with 1% PS and 0.2% BSA (Carrier solution). Cells were washed with PBS and treated with biotinylated secondary antibody (BA-100, Vector Laboratories, CA) diluted in Carrier solution for 90 min. Subsequently, cultures were washed and incubated with VECTASTAIN Elite ABC-HRP kit (PK-6100, Vector Laboratories, CA) 1:100 in PBS for 90 min. This was followed by washing and incubation in diaminobenzidine (DAB)-H202 solution (D4293-50, Sigma) for 10 min.

Thereafter, cultures were washed three times in 0.1 M sodium cacodylate buffer (pH 7.2–7.4) followed by osmication with 1% osmium tetroxide in cacodylate buffer. Dehydration using ascending ethyl alcohol concentration steps was followed by two rinses in propylene oxide. The embedding medium was infiltrated by first immersing the sections in a 1:1 solution of propylene oxide and Epon followed by neat Epon, and final polymerization at 60°C. 60 nm sections (ultrathin) were cut and analyses with a JEM-2100Plus Transmission Electron Microscope at 200kV (Jeol). Images were acquired with the XAROSA CMOS camera (Emsis).

### Figure preparation and statistical analysis

All image analysis was performed on the raw data. For some representative images, the linear background correction was applied equally for the groups within experiment. Figures were created using Adobe Illustrator software.

GraphPad Prism was used for statistical analysis. All data are represented as mean ± s.e.m. The data were tested with Grubb’s outlier test (α=0.05). To check for normal distribution the Shapiro-Wilk test was used. One-way ANOVA was followed by Tukey’s multiple comparison test. A nonparametric Kruskal-Wallis test was carried out for non-normally distributed data sets. For comparison between normally distributed data from two groups, a two-tailed, unpaired t-test was carried out. For comparison between non-normally distributed data of two groups, a two-tailed Mann-Whitney test was carried out. 2-way ANOVA followed by Tukey’s multiple comparison test was used to compare two variances (E-64 and Aβ3(pE)-42 oligomer treatment). P-values were considered as following: P > 0.05 = ns; ≤ 0.05 = *; ≤ 0.01 = **; ≤ 0.001 = ***; ≤ 0.0001 = ****. The type of statistical test used for each experiment, significance levels, the n number definition, and the look-up table (LUT) are reported in the corresponding figure legends.

## Results

### Aβ3(pE)-42 oligomers gradually accumulate in astrocytes

We have previously demonstrated that Aβ3(pE)-42 oligomer accumulations form within astrocytic but not neuronal cells (Grochowska *et al*., 2017). However, it remained unclear whether Aβ3(pE)-42 oligomers gradually accumulate in astrocytes or rather aggregate in the extracellular space prior to astrocytic uptake. To address this question, we treated primary astrocytic cultures with a previously characterized Aβ3(pE)-42 oligomeric preparation (Grochowska *et al*., 2017); Fig. S1A) for 0.5, 1, 4, 6, and 24 h (Fig. 1A-D). The samples were fixed and subsequently stained with antibodies detecting Aβ3(pE)-42 and GFAP, an astrocytic marker. We observed a gradual accumulation of Aβ3(pE)-42 oligomers as evidenced by the increase of mean intensity of Aβ3(pE)-42 deposits (Fig. 1A, B). Furthermore, the number of discrete Aβ3(pE)-42 signals decreased over time with a concomitant increase of Aβ3(pE)-42 area, indicating gradual accumulation of oligomers in bigger deposits (Fig. 1C, D). Of importance, when astrocytic cultures were treated with Aβ1-42 oligomers (Fig. S1B), we could not observe intracellular accumulation even after 72 h of treatment, indicating that the posttranslational Aβ modification required for accumulation in astrocytes (Fig. S1C).

**Fig. 1.**
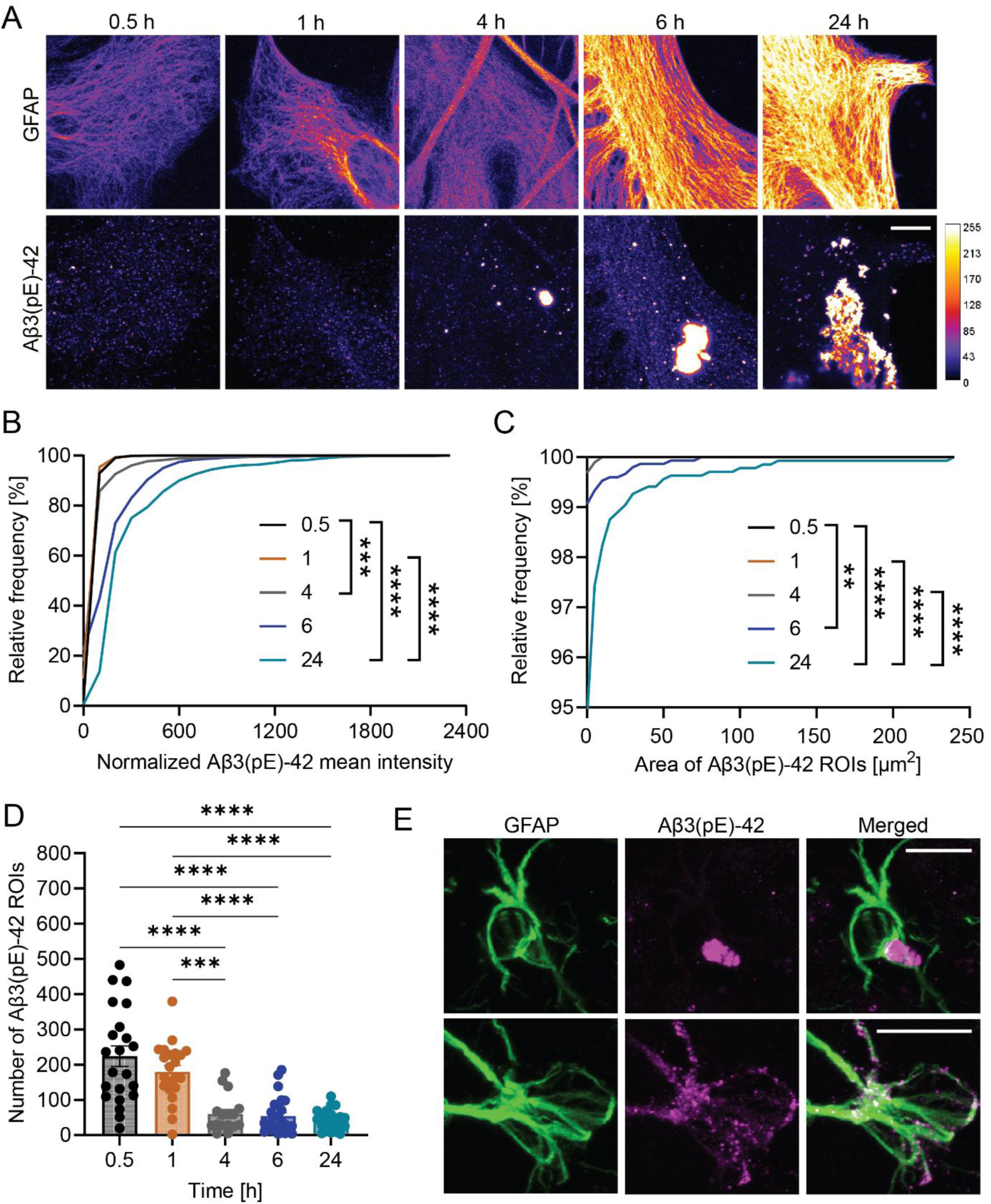
Aβ3(Aβ3(pE)-42)-42 oligomers gradually accumulate in primary murine astrocytes and glial cells of the TBA2.1 mice. A) Representative, confocal images of primary astrocytes treated with Aβ3(pE)-42 oligomers. Immunocytochemical detection of GFAP and Aβ3(pE)-42. The LUT shows the pixel intensities from 0 to 255. Scale bar, 10 μm. B) Relative, cumulative frequency distribution (in %) of normalized mean intensities of Aβ3(pE)-42 ROIs. Data is represented per ROI and normalized against the mean value of 0.5 h. The Aβ3(pE)-42 mean intensity increased with the duration of Aβ3(pE)-42 oligomer treatment. n = 16-25 cells from three independent experiments. C) Relative, cumulative frequency distribution (in %) of the area of Aβ3(pE)-42 ROIs. The area of ROIs increased with the duration of Aβ3(pE)-42 oligomer treatment. n = 16-25 cells from three independent experiments. D) Bar scatter plot of the number of Aβ3(pE)-42 ROIs per cell. The number of ROIs decreased with the duration of Aβ3(pE)-42 oligomer treatment. n = 16-25 cells from three independent experiments. Data are represented as mean ± s.e.m. E) Representative confocal images of brain sections of TBA2.1 mice show Aβ3(pE)-42 uptake in astrocytes. Immunocytochemical detection of GFAP and Aβ3(pE)-42 with nuclear (DAPI) counterstain. Scale bar, 5 μm (B-D) *\*p*< 0.05, *\*\*p*< 0.01, *\*\*\*p*< 0.001, *\*\*\*\*p*< 0.0001 by (D) one-way ANOVA followed by Tukey’s multiple comparison test or (B, C) Kruskal-Wallis test.

To verify, whether the Aβ3(pE)-42 astrocytic uptake occurs *in vivo* we analyzed brain samples from 24 weeks old TBA2.1 mice (Fig. 1E). TBA2.1 mice overexpress Aβ3(pE)-42 and at this age are characterized by prominent amyloid deposits, impairment in long-term potentiation, progressive neuronal loss in the CA1 region of the hippocampus, astrogliosis, and activated microglia (Alexandru *et al*, 2011). Immunostaining of coronal brain sections with Aβ3(pE)-42-specific antibody and GFAP revealed prominent Aβ3(pE)-42 inclusions in CA1 astrocytes (Fig. 1E). Of note, we could also observe Aβ3(pE)-42-positive microglial cells labelled with Iba-1 (Fig. S1D).

### Aβ3(pE)-42 uptake disrupts astrocytic endolysosomal pathway

We next investigated in which cellular compartment Aβ3(pE)-42-positive clusters accumulate. The endocytic pathway is a system of vesicles in which endocytosed cargo is gradually sorted via early endosomes (EE), late endosomes (LE), to the catabolic organelles, the lysosomes (Luzio *et al*, 2014; Nixon & Cataldo, 2006; Saftig & Klumperman, 2009; Van Acker *et al*, 2019). Lysosomes serve as a degradative endpoint not only for endosomal but also for the autophagic pathway and lysosomal dysfunction is thought to drive AD pathology (Knopman *et al*., 2021; Nixon, 2020; Nixon & Cataldo, 2006).

To investigate whether Aβ3(pE)-42 oligomers enter the endolysosomal system we used two measurements: the overall percentage of Aβ3(pE)-42 ROIs positive for a given endolysosomal marker and its mean immunofluorescence signal intensity within the Aβ3(pE)-42 ROIs. The cells were fixed after 0.5, 4, and 24 h of treatment which represented distinct stages of Aβ3(pE)-42 accumulation (Fig. 1A-D).

We first performed immunostaining with an antibody directed against a marker of EE, Rab5 (Luzio *et al*., 2014; Saftig & Klumperman, 2009; Van Acker *et al*., 2019). We observed that more than 40% of Aβ3(pE)-42 is positive for Rab5 already after 30 min of treatment (Fig. 2A, B). We did not observe differences in colocalization between the tested time points, which could be explained by continuous uptake of Aβ3(pE)-42 oligomers present in the media and/or abnormal sorting of endolysosomal membranes (Fig. 2A, B). Interestingly, however, we observed a gradual increase of Rab5 mean intensity within Aβ3(pE)-42 ROIs which points to the gradual accumulation of EE membranes (Fig. 2A, C). In addition, in accordance with previous studies demonstrating Rab5 upregulation in human post-mortem brains of AD patients (Ginsberg *et al*, 2010), we could observe an increase in the number of discrete Rab5 ROIs after 24 h of treatment (Fig. S2A). Next, we overexpressed RFP-Rab7 fusion protein, which labels LE. While the association after 30 min of treatment was below 15%, we could find significant recruitment of RFP-Rab7 membranes at later time points following treatment (Fig. 2D, E). Furthermore, like for Rab5, we could observe a gradual increase of the RFP-Rab7 signal within Aβ3(pE)-42 ROIs (Fig. 2D, F). To further characterize the membranes associated with Aβ3(pE)-42, we expressed the LAMP1 protein fused to GFP, a marker for late endosomes and lysosomes (Saftig & Klumperman, 2009). Similarly, to RFP-Rab7, we could observe an increase in the recruitment of LAMP1-GFP-positive membranes to the Aβ3(pE)-42 ROIs (Fig. 2G-I).

**Fig. 2.**
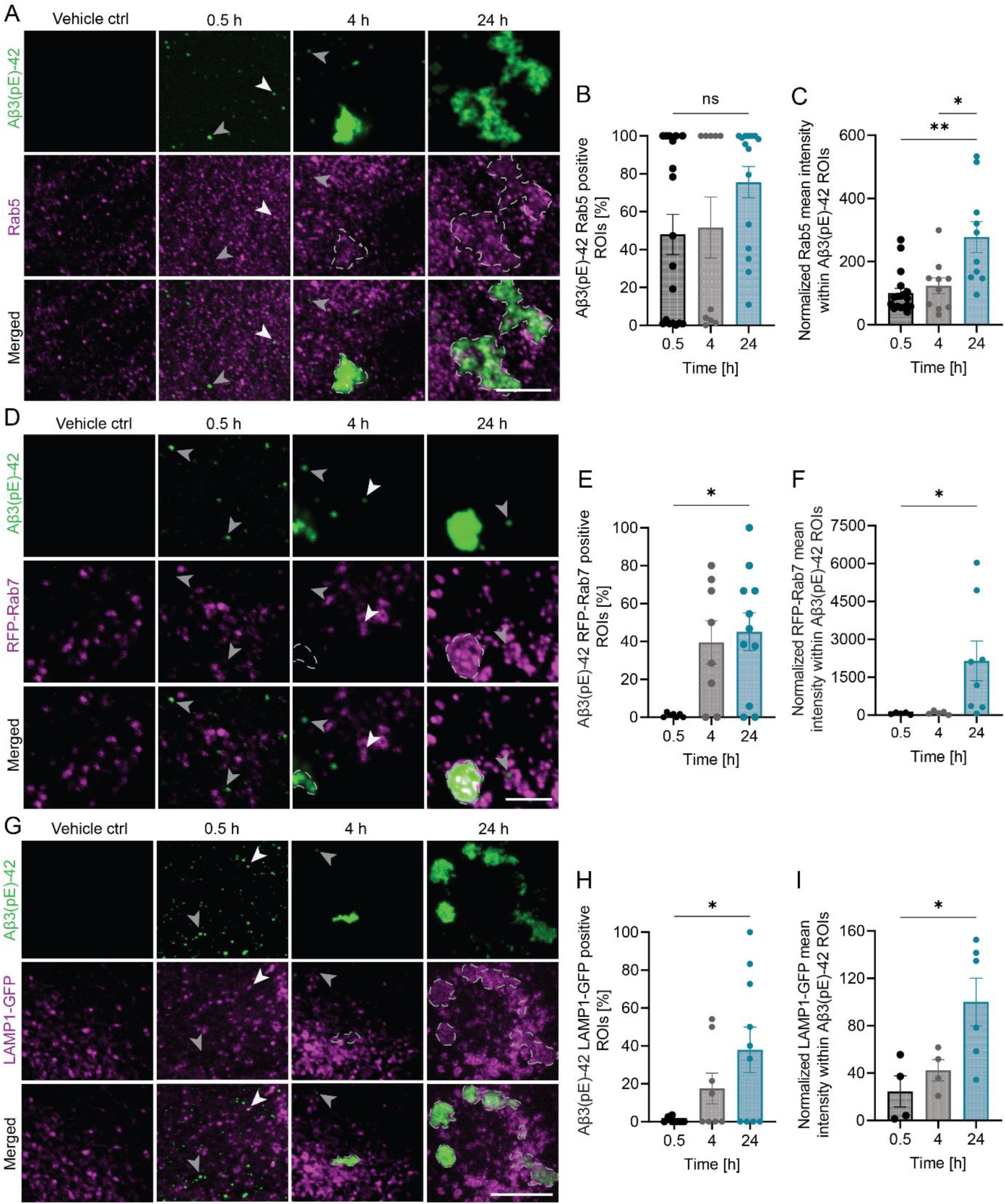
Uptake of Aβ3(pE)-42 oligomers disrupts the endolysosomal system in primary astrocytes. A) Representative, confocal images of primary astrocytes treated with Aβ3(pE)-42 oligomers. Immunocytochemical detection of Aβ3(pE)-42 and Rab5. White arrows indicate colocalization, grey arrows indicate no colocalization between Aβ3(pE)-42 and Rab5. Scale bar, 10 μm. B) Bar scatter plot of percentage of Aβ3(pE)-42 ROIs that colocalize with Rab5 per cell. n = 10-15 cells from three independent experiments. C) Bar scatter plot of the normalized Rab5 mean intensities within Aβ3(pE)-42 ROIs. Data is represented as average per cell and normalized against the mean value of 0.5 h. n = 10-15 cells from three independent experiments. D) Representative, confocal images of primary astrocytes treated with Aβ3(pE)-42 oligomers. Immunocytochemical detection of Aβ3(pE)-42 and overexpression of RFP-Rab7. White arrows indicate colocalization, grey arrows indicate no colocalization between Aβ3(pE)-42 and RFP-Rab7. Scale bar, 10 μm. E) Bar scatter plot of percentage of Aβ3(pE)-42 ROIs that colocalize with RFP-Rab7 per cell. n = 7-11 cells from two independent experiments. F) Bar scatter plot of the normalized RFP-Rab7 mean intensities within Aβ3(pE)-42 ROIs. Data is represented as average per cell and normalized against the mean value of 0.5 h. n = 7-11 cells from two independent experiments. G) Representative, confocal images of primary astrocytes treated with Aβ3(pE)-42 oligomers. Immunocytochemical detection of Aβ3(pE)-42 and overexpression of LAMP1-GFP. White arrows indicate colocalization, grey arrows indicate no colocalization between Aβ3(pE)-42 and LAMP1-GFP. Scale bar, 10 μm. H) Bar scatter plot of percentage of Aβ3(pE)-42 ROIs that colocalize with LAMP1-GFP per cell. n = 5-10 cells from two independent experiments. I) Bar scatter plot of the normalized LAMP1-GFP mean intensities within Aβ3(pE)-42 ROIs. Data is represented as average per cell and normalized against the mean value of 24 h. n = 8-10 cells from two independent experiments (B, C, E, F, H, I) Data are represented as mean ± s.e.m. *\*p*< 0.05, *\*\*p*< 0.01, *\*\*\*p*< 0.001, *\*\*\*\*p*< 0.0001 by (B, C, E, F) Kruskal-Wallis test or (H, I) by one-way ANOVA followed by Tukey’s multiple comparison test.

Autophagy is a lysosomal catabolic route in which cytoplasmic protein aggregates are sequestered within a double-membraned autophagosome (Nixon, 2020; Nixon & Cataldo, 2006; Van Acker *et al*., 2019). To investigate whether Aβ3(pE)-42 localizes to autophagosomes, we performed immunostainings with antibodies detecting LC3, a canonical autophagy marker (Ohsumi, 2014). We could not observe any significant recruitment of LC3 to the Aβ3(pE)-42 deposits (Fig. S2B-D). Furthermore, although the increase of autophagic flux induced by cellular starvation led to LC3 upregulation (Fig. S2B, E; (Ohsumi, 2014), we could not observe any influence on the area or number of Aβ3(pE)-42 ROIs or any change of its association with LC3 (Fig. S3B-G), indicating that this pathway does not significantly contribute to catabolism of Aβ3(pE)-42 (Fig. S2B, E, F).

Collectively, these data point to the gradual disruption of endolysosomal membrane sorting induced by the accumulation of Aβ3(pE)-42 oligomers.

### Aβ3(pE)-42 accumulation induces formation of heteromorphous vacuolar structures

We next tried to gather more information about the interaction of Aβ3(pE)-42 with the membranes of the endolysosomal system. To this end we analyzed astrocytic cells treated for 24h with Aβ3(pE)-42 oligomers using high-resolution transmission electron microscopy. The samples were stained with antibody against Aβ3(pE)-42 detected with diaminobenzidine (DAB). We identified single-membrane, heteromorphous vacuoles positive for Aβ3(pE)-42 deposits surrounded by smaller vesicles with Aβ3(pE)-42-positive membranes (Fig. 3A). The formation of large intracellular vacuoles with accumulations that vary in electron density are characteristic for lysosomal storage disorders (LSDs) and are associated with disrupted lysosomal degradation (Di Malta *et al*, 2012). Immunolabelling with an antibody detecting Cathepsin D showed that vacuoles from Aβ3(pE)-42 oligomer-treated cultures were also positive for this lysosomal enzyme (Fig. 3B). These results indicate the lysosomal nature of these vacuoles and corroborate lysosomal deficiency after stimulation. We also observed small Cathepsin D-positive vesicles that surround large membrane bound organelles with Cathepsin D positive storage material (Fig. 3C), strongly resembling the structures stained with antibody detecting Aβ3(pE)-42 (Fig. 3A). In nonstimulated control astrocytes Cathepsin D labelled-lysosomes were of typical rounded shape and homogenously distributed electron dense material (Fig. 3D).

**Fig. 3.**
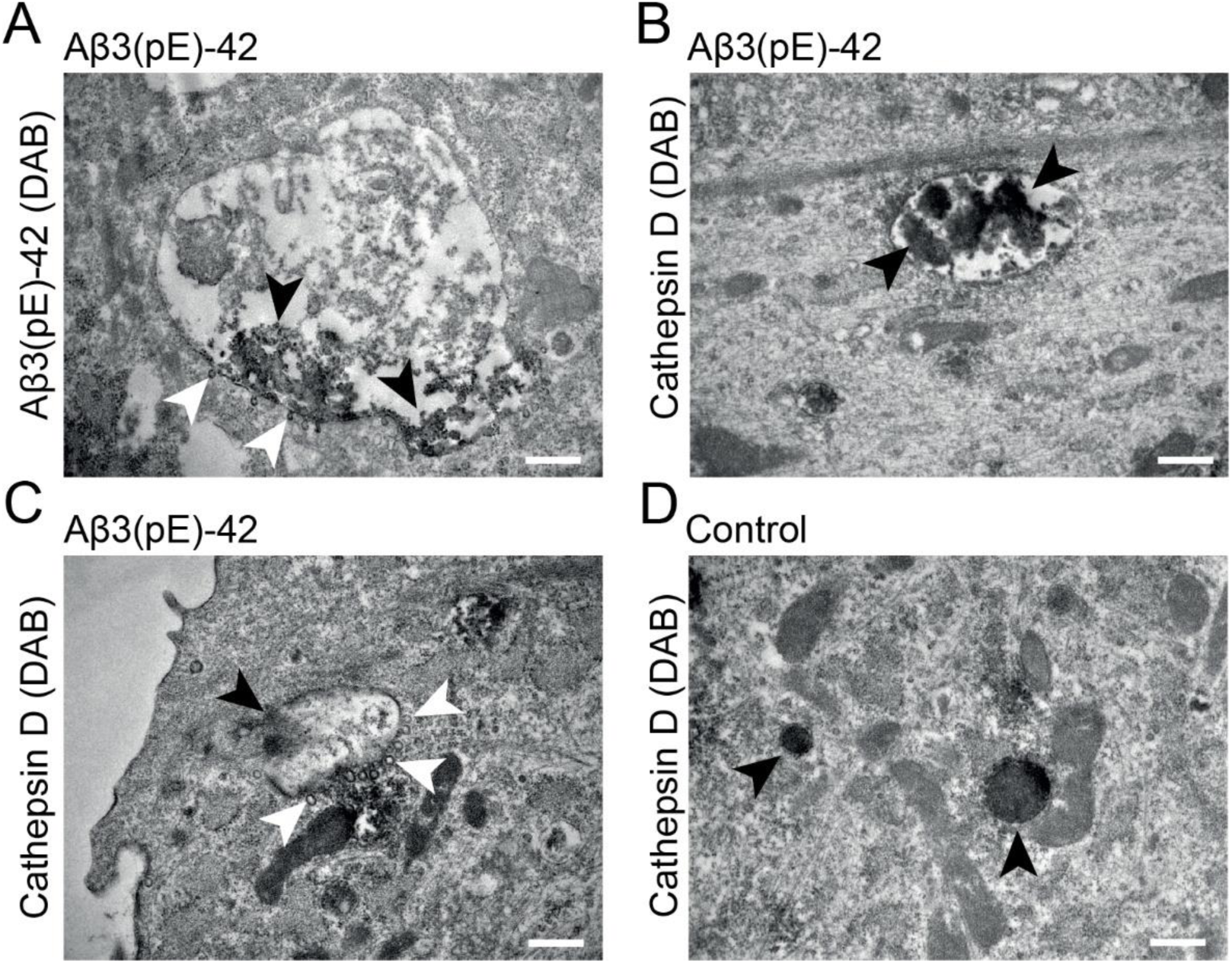
Aβ3(pE)-42 accumulate in heteromorphous vacuolar structures. A) Representative electron micrographs of primary astrocytes treated with Aβ3(pE)-42 oligomers for 24h stained with antibody against Aβ3(pE)-42 (DAB counterstain). Black arrows indicate Aβ3(pE)-42 deposits in single-membrane, hetermorphous vacuoles. White arrows indicate Aβ3(pE)-42 positive small vesicles surrounding large membrane bound organelles. Scale bar, 500 nm. B) Representative electron micrographs of primary astrocytes treated with Aβ3(pE)-42 oligomers. DAB-counterstain with antibody against Cathepsin D. Black arrow indicate heteromorphous vacuoles positive for Cathepsin D. Scale bar, 500 nm. C) Representative electron micrographs of primary astrocytes treated with Aβ3(pE)-42 oligomers. DAB-counterstain with antibody against Cathepsin D. Black arrow indicates heteromorphous vacuolar structure surrounded by smaller vesicles (white arrows) resembling structures positive for Aβ3(pE)-42. Scale bar, 500 nm. D) Representative electron micrographs of untreated primary astrocytes stained with antibody against Cathepsin D revealed regular, vesicular structures homogenously filled with Cathepsin D (black arrows). Scale bar, 500 nm.

### Aβ3(pE)-42 accumulation disrupts astrocytic lysosomal signaling

We therefore next wondered whether impairment in Aβ3(pE)-42 catabolism, disruption in endolysosomal membrane sorting and abnormal lysosomal morphology leads to the impairment of astrocytic lysosomal function. Lysosomal catabolism depends on low intravesicular pH that enables the activity of lysosomal proteases (Saftig & Klumperman, 2009). Therefore, to specifically label active compartments, we performed live imaging experiments with the SiR-lysosome dye, which binds to the active lysosomal protease, Cathepsin D (Lukinavičius *et al*, 2016). We used astrocytes incubated with fluorescently labeled Aβ3(pE)-42 oligomeric preparation (FAM-Aβ3(pE)-42; Fig. S3A). Lysosomes are highly motile organelles, and their active transport is essential for maintaining degradative functions (Lie & Nixon, 2019; Potokar *et al*, 2010). Indeed, in the control group, we could observe numerous SiR-lysosome-positive compartments of high motility (Fig. 4A-C). The incubation with FAM-Aβ3(pE)-42 for 24 h induced significant impairment of lysosomal velocity compared to vehicle-treated control (Fig. 4A-C). In line with the EM experiments (Fig. 3A, B), we observed colocalization of SiR-lysosome with small FAM-Aβ3(pE)-42 deposits (Fig. 4A, D). Furthermore, the low degree of labelling within bigger FAM-Aβ3(pE)-42 deposits indicated decreased catabolic activity (Fig. 4A-C).

**Fig. 4.**
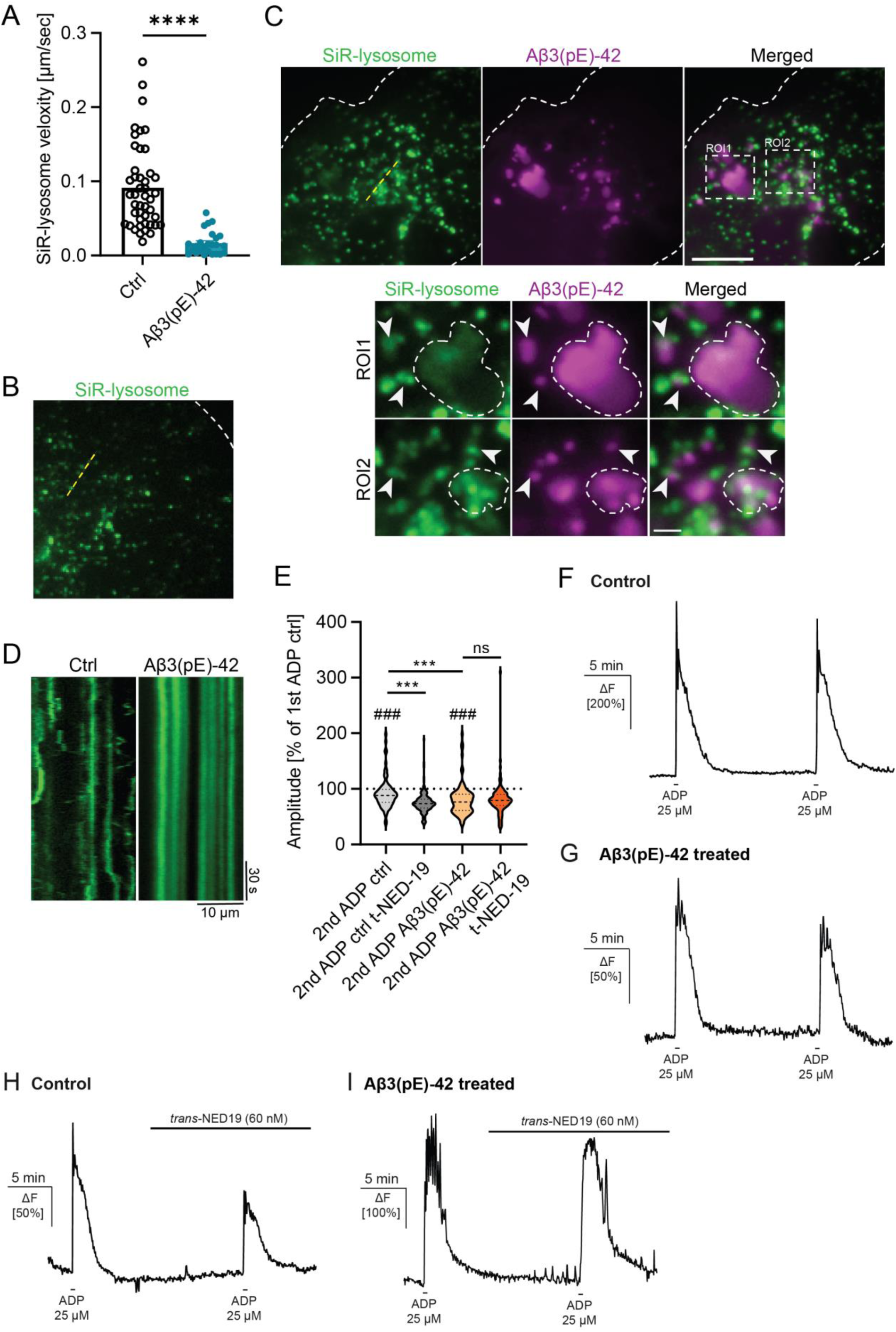
Aβ3(pE)-42 disrupts physiological lysosomal signaling. A) Bar scatter plot of the average of active lysosomal velocity per cell in non-treated and FAM-Aβ3(pE)-42 oligomers treated primary astrocytes. n = 25-45 cells from 2 independent experiments. B) Representative, maximum projection of live cell image streams of active lysosomes visualized with SiR-lysosome labelled Cathepsin D in FAM-Aβ3(pE)-42 oligomers treated primary astrocytes. C) Representative, maximum projection of live cell image stream of active lysosomes visualized with SiR-lysosome labelled Cathepsin D in FAM-Aβ (pE)-42 oligomers treated primary astrocytes. Scale bar, 10 µm. White arrows indicate colocalizations in ROI1 and ROI2. ROI1 shows the bigger FAM-Aβ3(pE)-42 deposits were vastly devoid of the active lysosomal proteases. ROI2 shows colocalization of SiR-lysosome with small FAM-Aβ3(pE)-42 deposits. Scale bar, 2 µm. D) Representative, kymographs of Cathepsin D labelled with SiR-lysosome from streams with a total acquisition time of 3 min (1 Hz acquisition frequency) reveal a significant decrease of SiR-lysosome velocity in FAM-Aβ(pE)-42 oligomers treated cells. Scale bar, 30 sec/ 10 µm. E-I) Aβ3(pE)-42 treatment disrupts lysosomal calcium signaling. (E) ADP-induced calcium signaling in primary astrocytes under control conditions is significantly reduced upon repetitive ADP application. The data are normalized to the mean of first ADP-induced calcium response (100% marked by dotted line). Application of *t*-Ned-19 significantly reduced the ADP-induced calcium response in control but not Aβ3(pE)-42-treated astrocytes. Data are represented as violin plots. n = 84-90 cells (control groups) 164 cells from 3 independent experiments. # denotes statistical comparisons of calcium responses between 1^st^ and 2^nd^ ADP application. * denotes comparisons of calcium responses after 2^nd^ ADP application with and without *r*-Ned-19. (F-I) Normalized mean amplitudes of the 1^st^ and 2^nd^ ADP-induced calcium responses in primary astrocytes in (F) control conditions, (G) Aβ3(pE)-42 oligomer-treated astrocytic cultures and (H, I) both groups pre-treated with *t-*Ned-19. Data are represented as mean ± s.e.m. *\*p*< 0.05, *\*\*p*< 0.01, *\*\*\*p*< 0.001, *\*\*\*\*p*< 0.0001 by (A, E - #) Mann-Whitney-U test and (E - *) one-way ANOVA followed by Kruskal-Wallis test.

Nicotinic acid adenine dinucleotide phosphate (NAADP)-dependent release of lysosomal Ca^2+^ is linked to lysosome deacidification or oxidative stress conditions (Lie & Nixon, 2019). Since abnormal Ca^2+^ signaling is a hallmark of astrocytic deregulation in neurodegenerative disorders, especially in AD (Lohr, 2023; Rodríguez-Arellano *et al*., 2016), we next looked at the astrocytic lysosomal Ca^2+^ signaling upon Aβ3(pE)-42 oligomer treatment. Aβ3(pE)-42 oligomer-treated astrocytic cultures as well as untreated control cultures were loaded with the chemical calcium indicator Rhod-2 AM, to visualize changes in cytosolic Ca^2+^ concentrations. To induce Ca^2+^ transients, we utilized the purinergic receptor agonist ADP, which activates P2Y1-receptor and the downstream Gq-dependent pathway *in-vitro* and *in-vivo* (Delekate *et al*, 2014b; Doengi *et al*, 2008; Fumagalli *et al*, 2003). ADP application induced robust Ca^2+^ transients with a mean amplitude of 193.93 ± 8.57 ΔF in control astrocytes and 198.43 ± 9.61 ΔF in Aβ3(pE)-42 oligomer-treated astrocytes (Fig 4F). To test for agonist-induced signalling cascade desensitization, we performed rundown experiment. ADP-induced Ca^2+^ transients showed a decrease in mean amplitude to 91.71 ± 4.13% of the 1^st^ ADP application in control astrocytes (Fig. 4E, F). In Aβ3(pE)-42 oligomer-treated astrocytes the second ADP application showed an even stronger decrease in amplitude to 80.35 ± 3.91% of the 1^st^ ADP application (Fig. 4E). The statistical comparison of the amplitudes of the 2^nd^ ADP-induced Ca^2+^ signals showed a highly significant difference between control and Aβ3(pE)-42 oligomer-treated astrocytes (Fig. 4E). To isolate the lysosome-dependent part of the ADP-induced Ca^2+^ transients in astrocytes we inhibited lysosomal Ca^2+^ release with *t*-Ned19 (60 nM), a structural analogue of NAADP (Morgan *et al*, 2011). As the previous results showed a decrease in amplitude with repetitive ADP applications (rundown) in both groups (Fig. 4E-G), we performed one control application of ADP (1^st^ ADP application) and compared the ADP application in the presence of *t*-Ned19 (2^nd^ ADP application) to the 2^nd^ ADP application of the corresponding rundown experiment, i.e. in the absence of *t*-Ned19 (Fig. 4E, H, I). In control astrocytes, we observed a significant decrease in mean amplitude in the presence of *t*-Ned19 compared to control (Fig. 4E, F, H), suggesting that around 17% of the Ca^2+^ response originates from the lysosomes (Fig. 4E, F, H). However, ADP-induced Ca^2+^ transients in the presence of *t*-Ned19 in Aβ3(pE)-42 oligomer-treated astrocytes did not significantly differ from the 2^nd^ ADP application in the rundown experiment, indicating lack of lysosomal Ca^2+^ release (Fig. 4 E, G, I). Taken together, the results suggest impaired lysosomal Ca^2+^ signaling in astrocytes treated with Aβ3(pE)-42 oligomers.

### Aβ3(pE)-42 leads to the lysosomal stress response, GFAP upregulation, and enhanced NF-кB reactivity

Due to its hydrophobic properties, Aβ3(pE)-42 can form stable, ion-permeable pores (Gillman *et al*., 2014; Lee *et al*., 2014) which in high concentrations could induce lysosomal membrane rupture. Therefore, we performed immunostainings of Aβ3(pE)-42-treated astrocytes with an antibody detecting pRab10, a marker of lysosomal damage (Bonet-Ponce & Cookson, 2022; Yan *et al*, 2018). pRab10 is the part of the lysosome membrane repair mechanism and its recruitment to the organelle surface upon persistent lysosomal damage is one of the prominent pathological features of AD (Bonet-Ponce & Cookson, 2022; Yan *et al*., 2018). The immunostaining with the antibody detecting pRab10 revealed an almost 2-fold increase in pRab10 immunoreactivity after 24 h of Aβ3(pE)-42 treatment (Fig. 5A, B). Furthermore, we could observe a prominent increase of the number of Aβ3(pE)-42 ROIs positive for pRab10 with more than 80% of positive ROIs after 24 h of treatment (Fig. 5A, C). We could not observe a significant increase in pRab10 mean fluorescence intensity within Aβ3(pE)-42 ROIs which could indicate limited activation of pRab10-dependent repair machinery associated with Aβ3(pE)-42-induced lysosomal impairment (Fig. 5A, D).

**Fig. 5.**
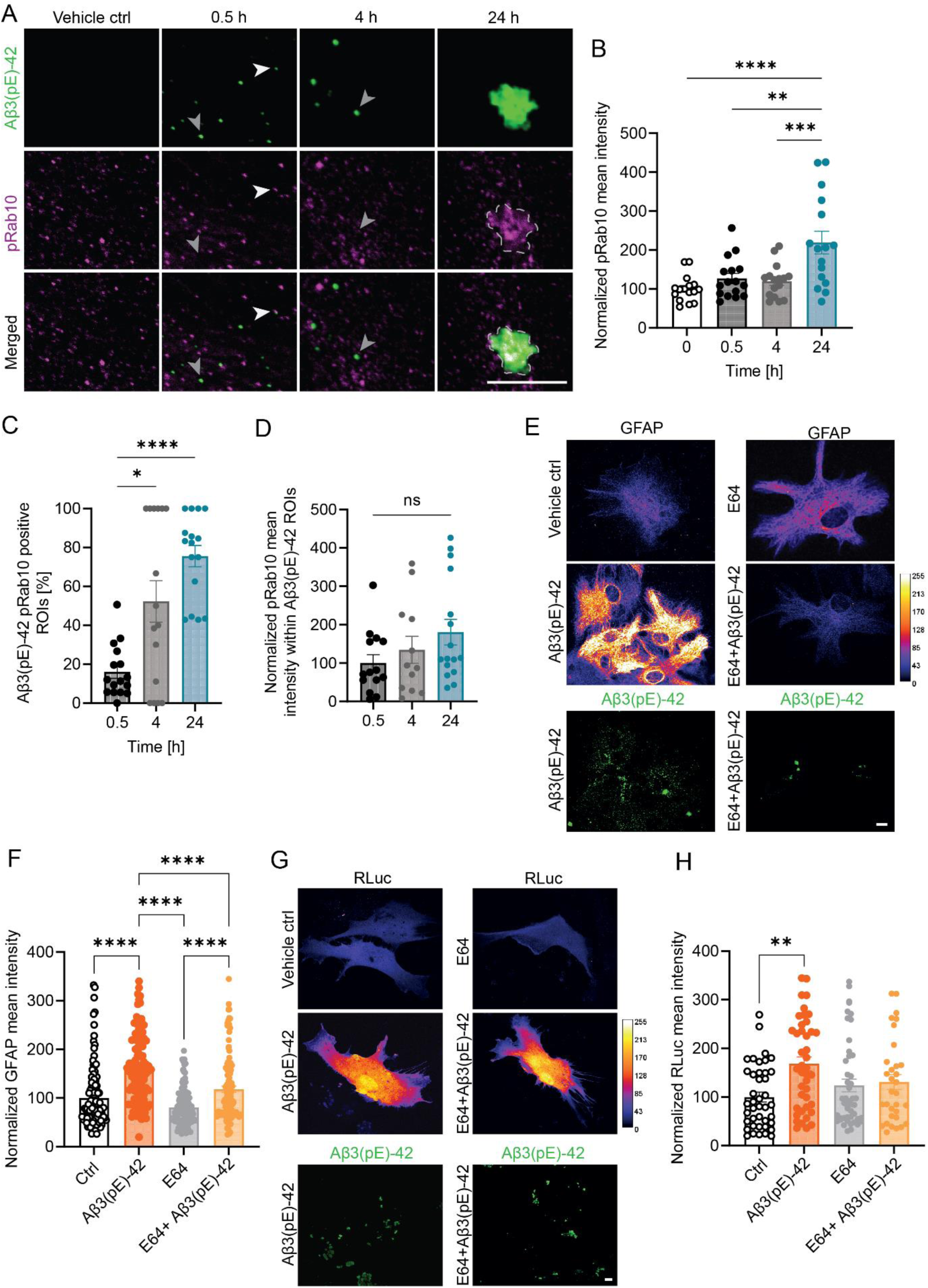
Aβ3(pE)-42 oligomers induce recruitment of pRab10 to the Aβ3(pE)-42-positive compartments, GFAP upregulation and NF-кB. A) Representative, confocal images of primary astrocytes treated with Aβ3(pE)-42 oligomers. Immunocytochemical detection of Aβ3(pE)-42 and pRab10. White arrows indicate colocalization, grey arrows indicate no colocalization between Aβ3(pE)-42 and pRab10. Scale bar, 10 μm. B) Bar scatter plot of the normalized mean intensities of pRab10 ROIs. Data is represented as average per cell and normalized against the mean value of vehicle control. n = 16 cells from three independent experiments. C) Bar scatter plot of percentage of Aβ3(pE)-42 ROIs that colocalize with pRab10 per cell. n = 16 cells from three independent experiments. D) Bar scatter plot of the normalized pRab10 mean intensities within Aβ3(pE)-42 ROIs. Data is represented as average per cell and normalized against the mean value of 0.5 h. n = 15-16 cells from three independent experiments. E) Representative, confocal images of primary astrocytes treated with Aβ3(pE)-42 oligomers with or without E-64. Immunocytochemical detection of Aβ3(pE)-42 and GFAP. Scale bar, 10 μm. F) Bar scatter plot of normalized GFAP mean immunoreactivity. n = 112-124 cells from two independent experiments. G) Representative, confocal images of primary astrocytes transfected with NF-кB GFP reporter treated with Aβ3(pE)-42 oligomers with or without E-64. Immunocytochemical detection of Aβ3(pE)-42. Scale bar, 10 μm. H) Bar scatter plot of normalized GFAP mean immunoreactivity. n = 32-47 cells from two independent experiments. (B-D, F, H) Data are represented as mean ± s.e.m. *\*p*< 0.05, *\*\*p*< 0.01, *\*\*\*p*< 0.001, *\*\*\*\*p*< 0.0001 by (C, D, F, H) Kruskal-Wallis test or (B) by one-way ANOVA followed by Tukey’s multiple comparison test.

It was previously suggested that Aβ3(pE)-42-induced lysosomal damage can lead to the lysosomal membrane permeabilization (LMP) and leakage of the proteases to the extracellular space (De Kimpe *et al*, 2013). This, in turn, was reported to drive the activation of neuroinflammatory pathways (Heneka *et al*, 2018; Wang *et al*, 2018). To verify, whether Aβ3(pE)-42 can lead to the activation of pro-inflammatory signaling, we looked at the upregulation of GFAP, a standard marker indicating astrocytic reactivity associated with astrocytic cluster characteristics for AD-associated astrocytes (Giovannoni & Quintana, 2020; Habib *et al*, 2020; Heneka *et al*., 2015; Liddelow & Barres, 2017). In line with previous report (Alexandru *et al*., 2011), we could confirm that TBA2.1 mice display GFAP upregulation in 2 hippocampal CA1 regions – stratum oriens and stratum radiatum (Fig. S4A-E). In addition, the treatment of primary astrocytes with Aβ3(pE)-42 oligomers for 0.5, 4, and 24 h revealed significant GFAP upregulation already after 4 h (Fig. S4F, G). On the other hand, we did not observe significant GFAP upregulation induced by Aβ1-42, even after 72 h of treatment (Fig. S3H, I). To verify whether abnormal activity of lysosomal proteases could drive GFAP upregulation, we treated cells with a broad lysosomal protease inhibitor, E-64 (Grinde, 1982). While the treatment with E-64 did not significantly change the levels of GFAP, the co-application of E-64 with Aβ3(pE)-42 oligomers abolished Aβ3(pE)-42-induced GFAP upregulation (Fig. 5E, G). Although GFAP upregulation is a proxy of astrocytic reactivity, alone it is insufficient to indicate the activation of pro-inflammatory pathway (Escartin *et al*, 2021; Liddelow & Barres, 2017). Therefore, we estimated the activity of p65/NF-кB, a transcription factor mediating pro-inflammatory cytokine expression (Heneka *et al*., 2018; Liddelow & Barres, 2017; Liu *et al*, 2017). To this end we transfected primary astrocytes with NF-кB expressing GFP under a high-affinity NF-кB promoter (Kuri *et al*, 2017). While we observed almost 70% increase in GFP fluorescence intensity relative to control, we could not observe NF-кB activation when the cells were treated with E-64 or Aβ3(pE)-42 and E-64 (Fig. 5G, H). Since NF-кB regulates apoptotic pathways (Liu *et al*., 2017) and LMP can result in cell death (Wang *et al*., 2018), we performed a luciferase-based, non-lytic cell viability assay (Fig. S4J). We could not detect any difference in the viability of astrocytic cultures followed by 24 h of treatment with Aβ3(pE)-42 oligomers (Fig. S4J). Collectively, these data indicate that Aβ3(pE)-42 astrocytic accumulations lead to LMP-associated induction of inflammatory signaling.

### The inhibition of lysosomal proteases rescues Aβ3(pE)-42-induced synapse loss

It was shown previously that astrocytes, but not neurons, take up Aβ3(pE)-42 and that this uptake leads to TNFα-dependent synaptic stripping (Grochowska *et al*., 2017). Since co-application of E-64 and Aβ3(pE)-42 oligomers decreased GFAP upregulation and NF-кB activation (Fig. 5), we tested whether this treatment would also rescue Aβ3(pE)-42-induced synapse loss. To this end we used primary, mix hippocampal cultures (DIV16), where neurons grow on top of astrocytic layer, and treated them with Aβ3(pE)-42 oligomers with and without E-64 (Fig. 6A, B). The quantification of apposing immunosignal of primary dendrites labeled with antibodies against synaptophysin and PSD-95 (pre- and post-synaptic marker respectively) revealed a significant decrease in the synaptic density in the case of Aβ3(pE)-42-treated, but not E-64 and Aβ3(pE)-42-treated neurons (Fig. 6A, B).

**Fig. 6.**
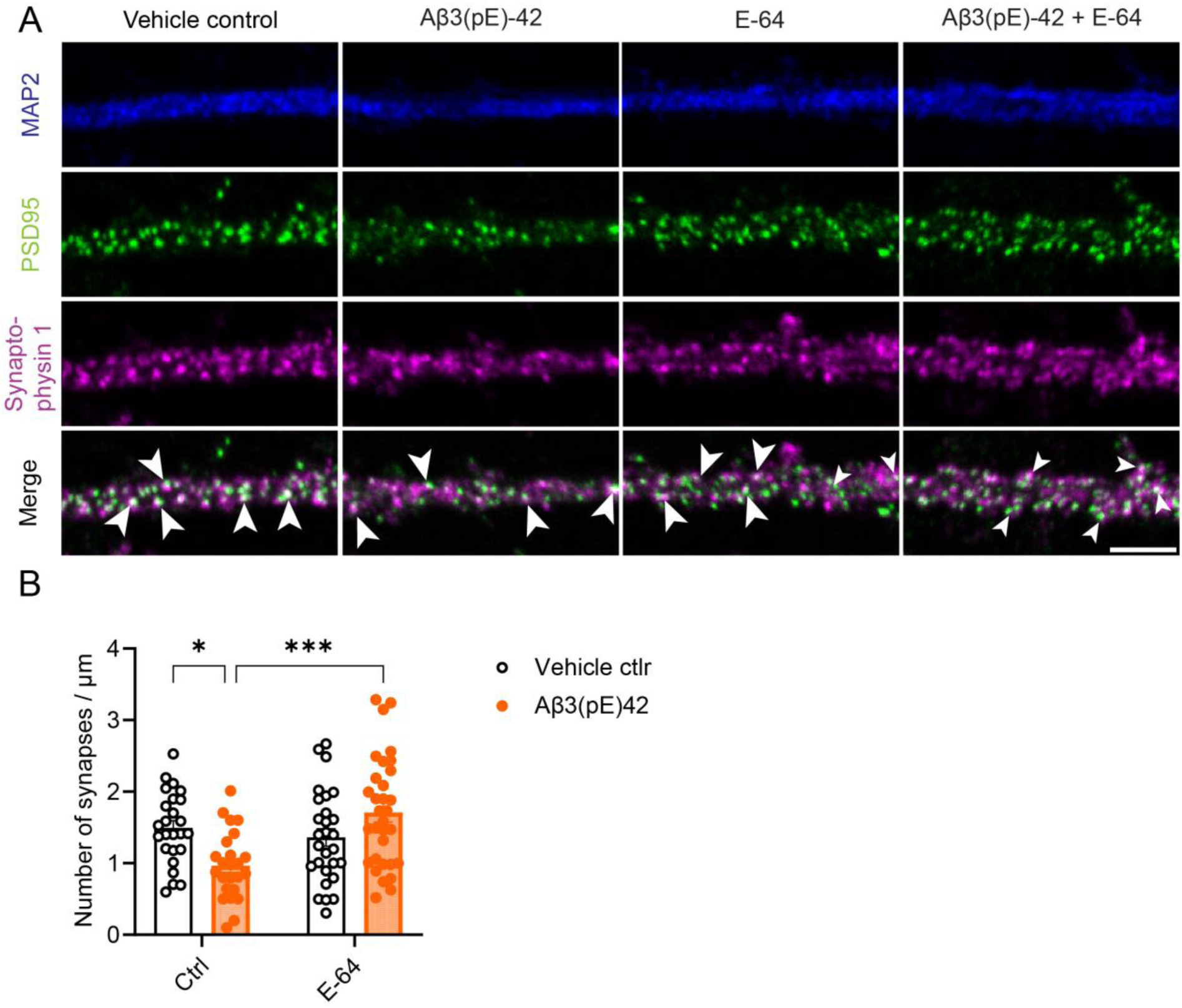
Co-application of E-64 rescues Aβ3(pE)-42-induced synapse loss in primary hippocampal neurons. A) Representative confocal images of hippocampal neurons treated with Aβ3(pE)-42 oligomers and E-64 for three days. Immunocytochemical detection of PSD95, Microtubule-associated protein 2 (MAP2), and synaptophysin 1. White arrows indicate synapses. Scale bar is 5 μm. B) Grouped bar scatter plot for the number of synapses per μm in untreated and E-64-treated neurons non-treated or treated with Aβ3(pE)-42 oligomers for three days. n = 17-20 cells from 2 independent cell cultures.

## Discussion

The impairment of Aβ clearance underlies the onset and progression of AD (De Strooper & Karran, 2016; Haass & Selkoe, 2022; Knopman *et al*., 2021; Selkoe & Hardy, 2016). It was previously demonstrated, that the most prevalent posttranslationally modified form of Aβ found in human brain, Aβ3(pE)-42, can be taken up by astrocytes which correlates with the release of TNFα, and, as a consequence, synaptic stripping (Grochowska *et al*., 2017; Harigaya *et al*., 2000; Jawhar *et al*., 2011; Kuo *et al*, 1997; Portelius *et al*., 2010; Saido *et al*., 1995). However, the mechanism underlying this process remained unclear. Here we demonstrate that Aβ3(pE)-42, but not Aβ1-42, gradually accumulates within the astrocytic endolysosomal system, leading to the impairment of lysosomal function and Ca^2+^ signaling, and lysosomal stress response. Furthermore, the inhibition of lysosomal proteases rescued the upregulation of GFAP and normalized activity of NF-кB as well as Aβ3(pE)-42-induced synapse loss. Taken together, this study supports the idea that posttranslational modifications of Aβ1-42 have significant implications for the mechanisms of impaired Aβ clearance underlying AD pathology. Furthermore, the data link the Aβ3(pE)-42-driven disruption of the endolysosomal system with induction of neuroinflammatory processes and subsequent synapse loss.

### Posttranslational modification of Aβ1-42 leads to clearance impairment

N-terminally truncated, pyroglutamylated amyloid-β peptides, including Aβ3(pE)-42, are the most prevalent modification detected in the brain of AD patients (Harigaya *et al*., 2000; Kuo *et al*., 1997; Portelius *et al*., 2010; Saido *et al*., 1995). These modifications constitute up to 50% of total Aβ in neuritic plaques (Harigaya *et al*., 2000; Kuo *et al*., 1997; Portelius *et al*., 2010; Saido *et al*., 1995). However, only very few studies have systematically compared Aβ species and related consequences for AD pathology. We did not observe astrocytic accumulation of Aβ1-42 even after 3 days of treatment with oligomers continuously present in the media. On the contrary, Aβ3(pE)-42 formed prominent accumulations already after 6 h of treatment. Since it was previously reported that Aβ1-42 is taken up by astrocytes and subsequently targeted to the lysosome for degradation (Heneka *et al*., 2015; Wyss-Coray *et al*, 2003), it is highly probable that the lack of visible accumulations is a result of efficient astrocytic clearance rather than differences in the uptake itself. Here we show than only the uptake of Aβ3(pE)-42 leads to the disruption of the endolysosomal system. This stunning difference between oligomeric species can be likely accounted for by biophysical properties of the N-terminal lactam ring of Aβ3(pE)-42 which imposes increased hydrophobicity, a tendency to form more stable β-sheet structures, faster aggregation, and higher resistance to degradation (De Kimpe *et al*., 2013; Jawhar *et al*., 2011; Schlenzig *et al*., 2009). Taking into account these properties, it is plausible that the enhanced Aβ3(pE)-42 concentration within the vesicles of the endolysosomal system combined with low pH could facilitate formation of β-sheet structures concomitantly decreasing degradation efficiency (Burdick *et al*, 1992). Interestingly, Aβ3(pE)-42 aggregates colocalize with LAMP1, a marker of the endolysosomal pathway, in the postmortem human brain tissue from patients suffering from AD (De Kimpe *et al*., 2013). This mechanism may be of particular relevance to AD pathology, especially since it was demonstrated that Aβ3(pE)-42 is able to seed toxic co-oligomers with Aβ1-42 and that co-oligomers form astrocytic accumulations in a similar manner as Aβ3(pE)-42 alone (Goldblatt *et al*, 2017; Grochowska *et al*., 2017; Nussbaum *et al*., 2012; Schlenzig *et al*., 2009).

### Aβ3(pE)-42 accumulation leads to disruption in endolysosomal sorting

Numerous mutations associated with both familial and late-onset AD indicate that the impairment in the endolysosomal system is central for defective Aβ clearance (Lee *et al*, 2010; Nixon, 2020; Van Acker *et al*., 2019). Although these genes are expressed across various cell types, the function of the astrocytic endolysosomal system is particularly important since astrocytes significantly contribute to Aβ clearance (De Strooper & Karran, 2016; Prasad & Rao, 2018; Selkoe & Hardy, 2016; Wong, 2020). Our results demonstrate that Aβ3(pE)-42 accumulation leads to formation of hybrid vacuoles positive for markers typical of various stages of the endolysosomal system – EE, LE, and mature lysosomes. These vacuoles can be divided into two groups based on their ultrastructure. Firstly, we observed small vesicles where Aβ3(pE)-42 localizes to the membranes rather than the lumen. This could be due to the peptide’s propensity to associate with lipid membranes (Gillman *et al*., 2014; Harigaya *et al*., 2000; Lee *et al*., 2014). These vesicles surrounded larger, heteromorphous structures resembling those observed in multiple sulfatase deficiency (MSD), a lysosomal storage disorder (LSD) characterized by progressive, severe neurodegeneration (Di Malta *et al*., 2012). Along these lines, early activation of astrocytes and GFAP upregulation is a hallmark of many LSDs, emphasizing the essential role of astrocytic endolysosomal disruption in driving neuroinflammation, neurodegeneration, and cognitive impairment (Bosch & Kielian, 2015; Wong, 2020).

### Aβ3(pE)-42 accumulation impairs lysosomal Ca^2+^ signals

The second finding of this study concerns the impairment in lysosomal physiological signaling. Astrocytes are the main trophic cells of the brain, and the impairment in their physiology may alter the trajectory of neurodegeneration, exacerbating synaptic impairment and neuronal loss (De Strooper & Karran, 2016; Di Malta *et al*., 2012; Haass & Selkoe, 2022; Patani *et al*., 2023; Rodríguez-Arellano *et al*., 2016). Ca^2+^ signals are key in mediating astrocyte-neuron and astrocyte-astrocyte communication (Nanclares *et al*, 2021; Semyanov *et al*, 2020; Zorec *et al*, 2012). Along these lines, it was reported that impairment in lysosomal function is causative for the disruption of ATP-mediated Ca^2+^ signals and glutathione (GSH) secretion and precedes neuronal loss in the model of Batten disease (Lange *et al*, 2018; Parviainen *et al*, 2017). In AD, Ca^2+^ signaling in astrocytes is upregulated (Lohr, 2023; Nanclares *et al*., 2021). Activated astrocytes in AD mouse models exhibit increased frequency of spontaneous Ca^2+^ transients and P2Y_1_-dependent Ca^2+^ hyperactivity that leads to neuronal network disfunction (Delekate *et al*, 2014a; Reichenbach *et al*, 2018; Takano *et al*, 2007). In APP/PS1 mice, Ca^2+^ signals synchronize and travel as Ca^2+^ waves over long distances uncoupled from neuronal activity (Kuchibhotla *et al*, 2009). However, which role lysosomal Ca^2+^ stores play in AD-mediated alterations of astrocytic Ca^2+^ signaling needs further investigation. In a mouse model of Huntington’s disease, Ca^2+^ release from astrocytic lysosomes fosters aggregation of huntingtin (mHtt), emphasizing the important role of lysosomal Ca^2+^ signaling in astrocytes in neurodegenerative diseases (Pereira *et al*, 2023).

### Disruption of the endolysosomal pathway links to induction of inflammatory signaling

Previous research on the role of astrocytes in neurodegeneration that underlie lysosomal storage disorders provides additional evidence for a link between the loss of physiological endolysosomal system function and changes in astrocytic cytokine expression (Lange *et al*., 2018; Parviainen *et al*., 2017; Sanyal *et al*, 2020). We therefore reasoned, that the role of Aβ3(pE)-42-induced, astrocyte-driven neurotoxicity may be not only be due to the loss of trophic functions but also due to the induction of synaptotoxic, pro-inflammatory signaling. Together with microglia, reactive astrocytes contribute to pro-inflammatory, neurotoxic signaling in AD by secreting neurotoxic agents, including pro-inflammatory cytokines (De Strooper & Karran, 2016; Heneka *et al*., 2015; Liddelow *et al*, 2017). Indeed, the loss of synapses induced by Aβ3(pE)-42 (Cho *et al*, 2023) depends on the release of TNFα from astrocytes (Grochowska *et al*., 2017). The current study suggests that Aβ3(pE)-42-induced lysosomal destabilization and lysosomal membrane permeabilization (LMP) are casual for the induction of pro-inflammatory response. In fact, leakage of lysosomal proteins drives both priming and activation of the nod-like receptor protein 3 (NLRP3) inflammasome, a mechanism relevant also in AD (Ditaranto *et al*, 2001; Halle *et al*, 2008; Hornung *et al*, 2008; Yang *et al*, 2019). An analogous mechanism was described in Parkinson’s disease (PD), a neurodegenerative disorder where disruption of LMP caused by pathological α-synuclein aggregates is linked to neuroinflammation and neuronal cell loss mediated by astrocytes (Booth *et al*, 2017; Jiang *et al*, 2017; Qiao *et al*, 2016). Qiao and colleagues demonstrated that a PD-associated mutation in the *Atp13a2* (Park9) gene, which encodes a lysosomal ATPase, leads to increased LMP in astrocytes (Qiao *et al*., 2016). As a result, Cathepsin B is released into the cytoplasm and subsequently induces NLRP3 inflammasome-driven IL1β production (Qiao *et al*., 2016). In this regard NFкB might be a key regulator of inflammation and astrocytic activation of NFкB and subsequent release of synaptotoxic signaling molecules is associated with many neurodegenerative conditions, including AD (González-Reyes *et al*, 2017; Kaltschmidt *et al*, 1997; Lian *et al*, 2015). In support of this notion, it has been shown that Aβ-activated astrocytes display significant upregulation of NFкB and subsequent release of the neurotoxic complement factor C3 which leads to alteration of dendritic morphology and synapse loss (Lian *et al*., 2015).

## Conclusions

In conclusion, these results highlight the importance of posttranslational modification of Aβ in the pathophysiology of AD. Furthermore, although astrocytes may not necessarily initiate the disease onset, the disruption of the astrocytic endolysosomal system and, as consequence, impairment in normal neuronal support functions and induction of inflammatory signaling, may plausibly significantly contribute to disease severity and progression. Therefore, these results prompt the need to further investigate the mechanism underlying LMP and induction of inflammation to fully exploit potential strategies preventing both the induction of inflammatory signaling and boosting astrocytic Aβ catabolism and related clearance processes. Furthermore, it will be important to deepen the understanding about the contribution of different glia types *in vivo*.

## Author contriubtions

KMG designed the study. KMG, MS, AB, CL wrote the first draft of the paper. SW, AS, AB, MS, KMG performed experiments. SW, AS, AB, MS, KMG analyzed data. SW, AS prepared the figures. KMG and CL acquired the funding. All authors read and commented on the manuscript.

## Acknowledgements (funding statement)

We gratefully acknowledge the professional technical assistance of C.Raithore and E. Szpotowicz. We would like to thank Dr. A.V. Failla from the UKE Microscopy Imaging Facility (DFG Research Infrastructure; RI_00489). The study was supported by Alzheimer Forschung Initiative e.V DE-20057p and CRC 1436 TPA02 to KMG, SFB 1328 (project number 335447717) to CL.

## Data availability statement

The data that support the findings of this study are available from the corresponding author upon reasonable request.

## Ethics approval statement

All animal experiments were carried out following the European Communities Council Directive (2010/63/EU) and approved by the local authorities.

