## Supporting-information for "Astrocytic uptake of posttranslationally modified amyloid-β leads to endolysosomal system disruption and induction of pro-inflammatory signaling"

### Supporting figures

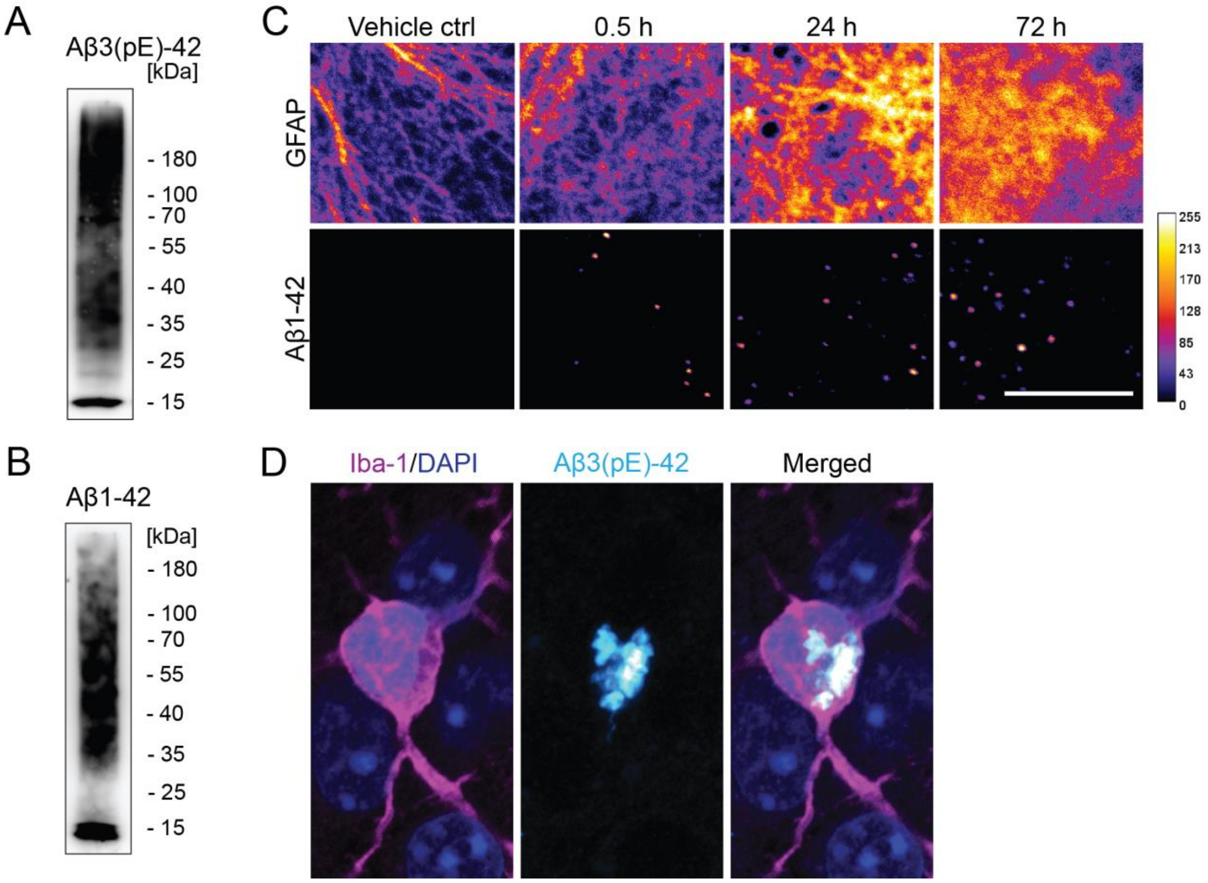


Fig. S1 – relates to Fig. 1

A) SDS-PAGE characterization of Aβ3(pE)-42 oligomeric preparation.

B) SDS-PAGE characterization of Aβ1-42 oligomeric preparation.

C) Representative, confocal images of primary astrocytes treated with Aβ1-42 oligomers show no formation of Aβ1-42 deposits within astrocytes. Immunocytochemical detection of Aβ1-42 and GFAP. LUT shows the pixel intensities from 0 to 255. Scale bar, 10 µm.

D) Representative confocal images of hippocampal cryosections of TBA2.1 mice show Aβ3(pE)-42 uptake in microglia cells. Immunocytochemical detection of Iba1 and Aβ3(pE)-42. Staining of cell nuclei with DAPI. Scale bar, 5 µm.

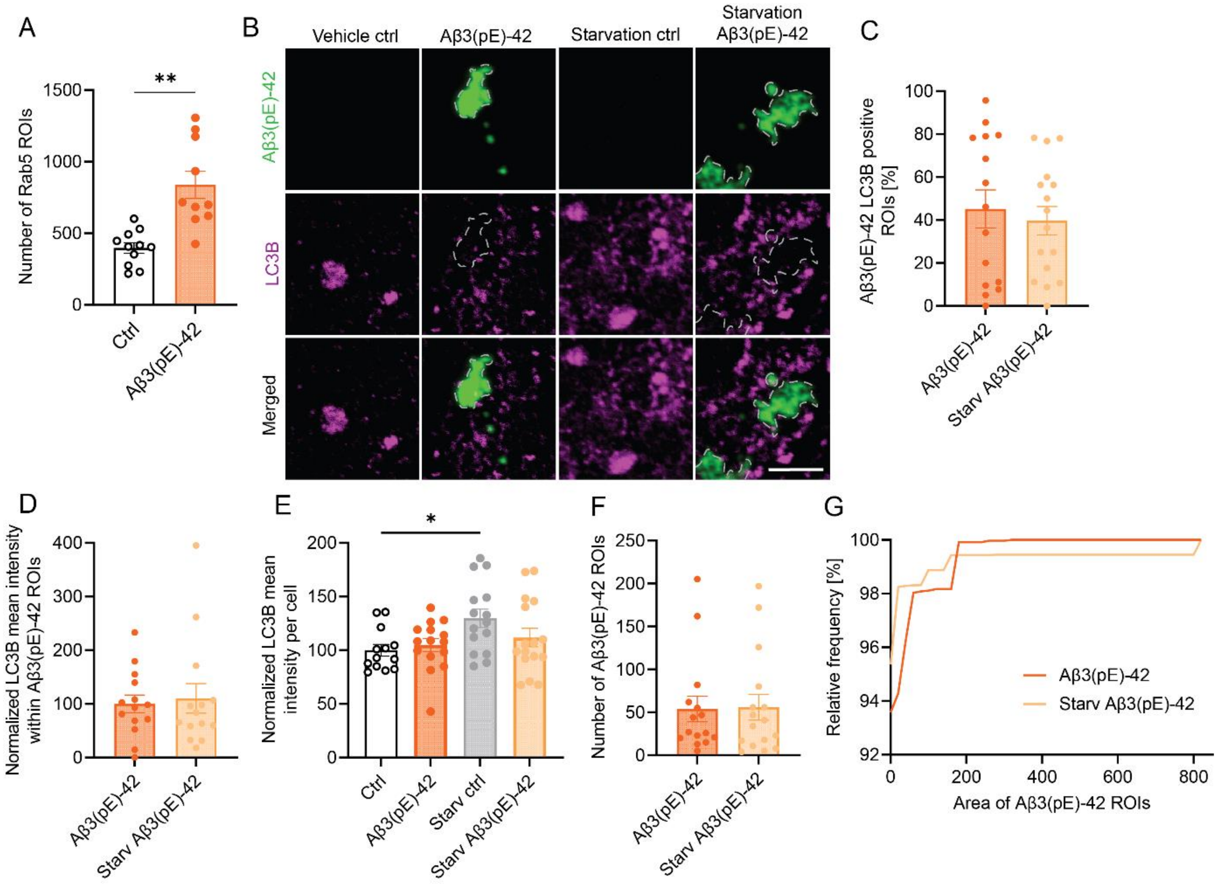


Fig. S2 – relates to Fig. 2

A) Bar scatter plot of the number of Rab5 ROIs per cell. The number of Rab5 ROIs increased with the duration of Aβ3(pE)-42 oligomer treatment. n = 10-18 cells from three independent cultures.

B) Representative, confocal images of primary astrocytes treated with Aβ3(pE)-42 oligomers. Treatment in serum-free media (starvation) induced upregulation of LC3 but no Aβ3(pE)-42 clearance. Immunocytochemical detection of Aβ3(pE)-42 and LC3B. Scale bar, 5 µm.

C) Bar scatter plot of percentage of Aβ3(pE)-42 ROIs that colocalize with LC3B per cell. n = 15-16 cells from one experiment.

D) Bar scatter plot of the normalized LC3B mean intensities within Aβ3(pE)-42 ROIs. Data is represented as average per cell and normalized against the mean value of Aβ3(pE)-42. n = 15-16 cells from one experiment.

E) Bar scatter plot of the normalized LC3B mean intensities. Starvation induced upregulation of LC3B in primary astrocytes. Data is represented as average per cell and normalized against the mean value of vehicle control. n = 15-16 cells from one experiment.

F) Bar scatter plot of the number of Aβ3(pE)-42 ROIs per cell. n = 15-16 cells from one experiment.

G) Relative, cumulative frequency distribution (in %) of the area of Aβ3(pE)-42 ROIs. The area of ROIs increased with the duration of Aβ3(pE)-42 oligomer treatment. n = 7 cells from one experiment.

(A, C-F) Data are represented as mean ± s.e.m. *$p< 0.05$, **$p< 0.01$, ***$p< 0.001$, ****$p< 0.0001$ by (A, E) one-way ANOVA followed by Tukey's multiple comparison test.

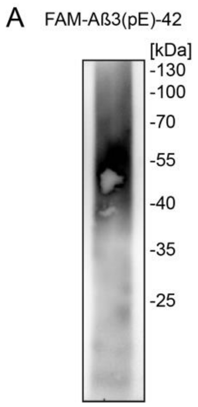


Fig. S3 – relates to Fig. 4
A) SDS-PAGE characterization of FAM-Aβ3(pE)-42 oligomeric preparation.

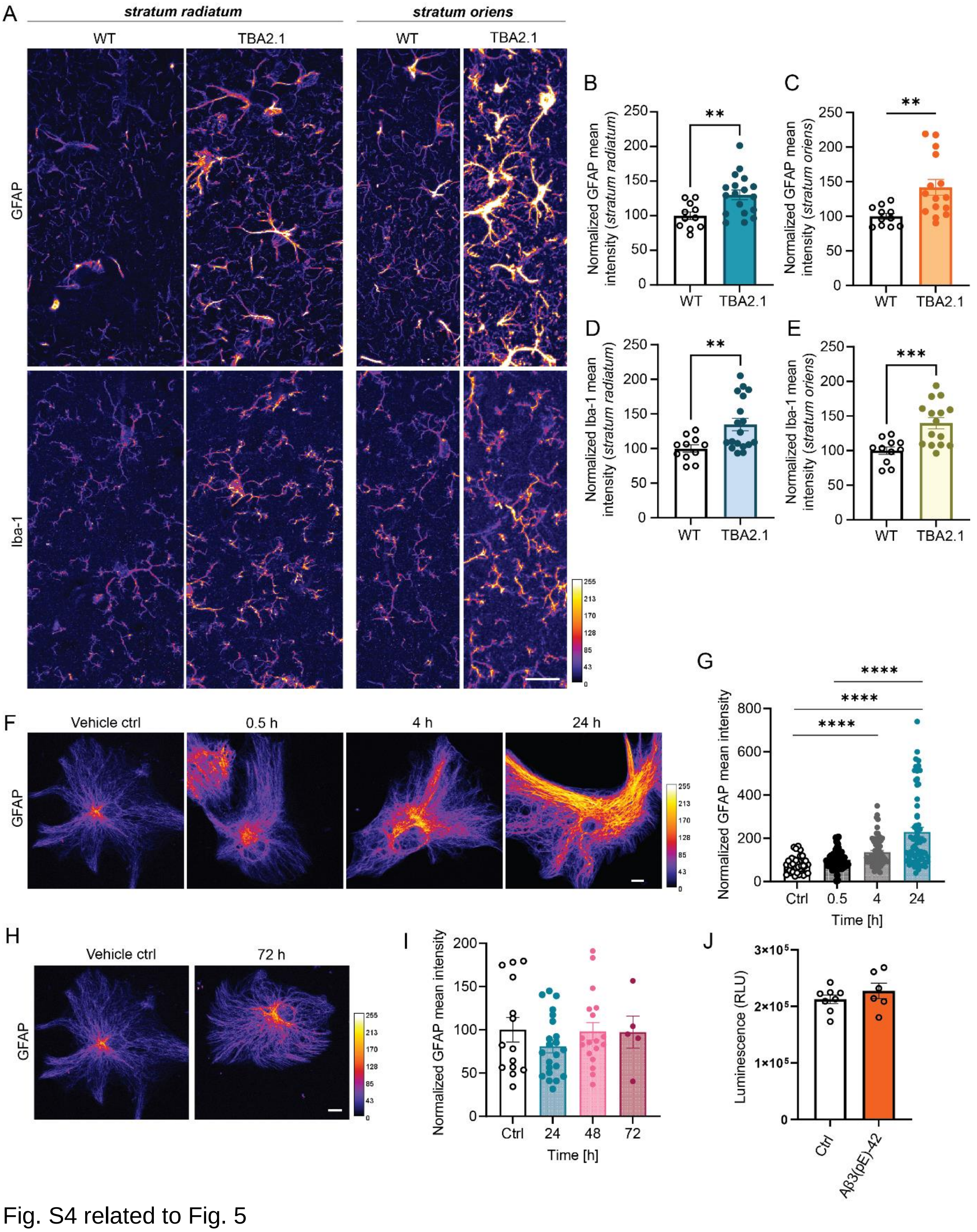


Fig. S4 related to Fig. 5

A) Representative, confocal images of primary astrocytes treated with Aβ3(pE)-42 oligomers. Immunocytochemical detection of GFAP, media without antibiotics. LUT shows the pixel intensities from 0 to 255. Scale bar, 10 μm.

B) Bar scatter plot of the normalized GFAP mean intensities per cell after Aβ3(pE)-42 treatment. Data is normalized against the mean value of 0.5 h. n = 40-64 cells from three independent experiments.

C) Representative confocal images of TBA2.1 hippocampal slices indicate upregulation of glial markers. Immunocytochemical detection of GFAP and Iba1 in stratum radiatum and stratum oriens. Scale bar, 20 μm.

D) Bar scatter plot of normalized GFAP mean intensity in the slices of WT and TBA2.1 mice in stratum radiatum. Data is normalized against the mean value of control. n = 11-18 slices from 3-5 animals.
E) Bar scatter plot of normalized GFAP mean intensity in the slices of WT and TBA2.1 mice in stratum oriens. Data is normalized against the mean value of control. n = 11-15 slices from 3-5 animals.
F) Bar scatter plot of normalized Iba1 mean intensity in the slices of WT and TBA2.1 mice in stratum radiatum. Data is normalized against the mean value of control. n = 11-18 slices from 3-5 animals.
G) Bar scatter plot of normalized Iba1 mean intensity in the slices of WT and TBA2.1 mice in stratum oriens. Data is normalized against the mean value of control. n = 11-15 slices from 3-5 animals.
H) Representative, confocal images of primary astrocytes treated with Aβ1-42 oligomers. Immunocytochemical detection of GFAP, media without antibiotics. LUT shows the pixel intensities from 0 to 255. Scale bar, 10 μm.
I) Bar scatter plot of the normalized GFAP mean intensities per cell after Aβ1-42 treatment, media without antibiotics. Data is normalized against the mean value of vehicle control. n = 5-22 cells from at least one experiments.
J) Relative luminescence (RLU) values corresponding to the substrate activated in living cells. n = 6-8 wells from one experiment.
(B, E, F, H, I) Data are represented as mean ± s.e.m. *$p$< 0.05, **$p$< 0.01, ***$p$< 0.001, ****$p$< 0.0001 by (B, H) Kruskal-Wallis test or (E, F) by Mann-Whitney test.

**Table S1. Antibodies, chemicals, kits, recombinant proteins, peptides, and software used in the study.**

| REAGENT or RESOURCE | SOURCE | IDENTIFIER |
|---|---|---|
| **Antibodies** | | |
| Rabbit anti-Aβ-pE3 | Synaptic Systems | Cat.: #218 003; RRID: AB_2056424 |
| Mouse anti-Aβ (4G8) | BioLegend | Cat.: #800701; RRID: AB_2564633 |
| Guinea pig anti-GFAP | Synaptic Systems | Cat.: #173 004; RRID: AB_10641162 |
| Rabbit anti-GFAP | Sigma-Aldrich | Cat.: #G-9269; RRID: AB_477035 |
| Guinea pig anti-IBA1 | Synaptic Systems | Cat.: #234004; RRID: AB_2493179 |
| Mouse anti-Rab5A (E6N8S) | Cell Signaling Technologies | Cat.: #46449; RRID: AB_2799303 |
| Rabbit anti-LC3B | Abcam | Cat.: #ab48394; RRID: AB_881433 |
| Rabbit anti-pRAB10 (phospho T73) | Abcam | Cat.: #ab241060; RRID: AB_2884876 |
| Rabbit anti-Cathepsin D | Abcam | Cat.: #ab72915 RRID: AB_2040714 |
| Guinea pig anti-MAP2 | Synaptic Systems | Cat.: #188 004; RRID: AB_2138181 |
| Rabbit anti-PSD95 | Nanotag Biotechnologies | Cat.: #N3783; RRID: N/A |
| Mouse anti-Synaptophysin 1 | Synaptic Systems | Cat.: #101 011; RRID: AB_887824 |
| Anti-guinea pig Alexa Fluor 405 | Abcam | Cat.: #ab175678; RRID: AB_2827755 |
| Anti-guinea pig Alexa Fluor 488 | Thermo Fisher Scientific | Cat.: #A-11073; RRID: AB_2534117 |

| | | |
|---|---|---|
| Anti-mouse Alexa Fluor 488 | Thermo Fisher Scientific | Cat.: #A-11001; RRID: AB_2534069 |
| Anti-mouse Alexa Fluor 568 | Thermo Fisher Scientific | Cat.: #A-11031; RRID: RRID: AB_144696 |
| Anti-mouse Alexa Fluor 647 | Thermo Fisher Scientific | Cat.: #A-31571; RRID: AB_162542 |
| Anti-mouse-IgG-HRP | Dianova | Cat.: #115-035-146; RRID: AB_2307392 |
| Anti-rabbit Alexa Fluor 488 | Thermo Fisher Scientific | Cat.: #A-11034; RRID: AB_2576217 |
| Anti-rabbit Alexa Fluor 568 | Thermo Fisher Scientific | Cat.: #A-11036; RRID: AB_10563566 |
| Anti-rabbit Alexa Fluor 647 | Thermo Fisher Scientific | Cat.: #A-32733; RRID:AB_162542 |
| Anti-rabbit-IgG-HRP | Dianova | Cat.: #111-035-114; RRID: AB_2337938 |
| Anti-Rabbit IgG Biotinylated | Vector Laboratories | Cat: #BA-100 RRID:AB_2313606 |
| **Bacterial and virus strains** | | |
| *E. coli* XL10Gold | Agilent | Cat.: #200314 |
| **Chemicals, media, and peptides** | | |
| Lipofectamine 2000 | ThermoFisher Scientific | Cat.: #11668-019 |
| SiR-Lysosome | Spirochrome | Cat.: #SC012 |
| E-64 protease inhibitor | Sigma-Aldrich | Cat.: #E3132-1MG |
| [Pyr3]-beta-Amyloid (3-42) | Anaspec | Cat.: #AS-29907 |
| Beta-Amyloid (1-42) | Anaspec | Cat.: #AS-20276 |
| [Pyr3]-beta-Amyloid (3-42)FAM labelled | Eurogentec | Custom made |
| Deoxyribonuclease I | Roche | Cat.: #11284934001 |
| L-glutamine | Thermo Fisher Scientific | Cat.: #25030149 |
| Hank's Balanced Salt Solution | Sigma | Cat.: #H9269-500ML |
| Neurobasal medium | Gibco | Cat.: #21103049 |
| F12 medium | Thermo Fisher Scientific | Cat.: #21765029 |
| Fetal Bovine Serum | Thermo Fisher Scientific | Cat.: #10082147 |
| Diaminobenzidine (DAB)-H202 solution | Sigma-Aldrich | Cat: #D4293-50 |
| Deoxyribonuclease I | Roche | Cat.: #11284934001 |
| Dimethyl sulfoxide | Sigma-Aldrich | Cat.: #276855 |
| Rhod-2 AM | Abcam | Cat.: #ab142780 |
| *t*-NED19 | Bio-techne | Cat.: #3954 |
| Adenosindiphosphate (ADP) | AppliChem | Cat.: #A0948 |
| **Critical commercial assays** | | |
| RealTime-Glo MT Cell Viability Assay | Promega | Cat.: #G9711 |
| VECTASTAIN Elite ABC-HRP Kit | Vector Laboratories | Cat: #PK-6100 |
| NucleoBond Xtra Midi EF | Machery-Nagel GmbH | Cat.: #740420.50 |
| NucleoSpin Gel and PCR Clean-up | Machery-Nagel GmbH | Cat.: #12870256 |
| PierceTM Bradford Assay Kit | Thermo Fisher Scientific | Cat.: #23200 |
| RealTime-Glo MT Cell Viability Assay | Promega | Cat.: #G9711 |
| **Experimental models: Organisms/strains** | | |
| Rat: Wistar | Janvier | RjHan:WI |

| | | |
|---|---|---|
| Mouse: B6;129-Gt(ROSA)26Sor$^{tm1(CAG-cas9*,-EGFP)Fezh}$/J | Jackson laboratory, (Platt *et al*, 2014) | Cat.: #024857 |
| Mouse: JAX C57BL/6 | Jackson laboratory | Cat.: #000664 |
| Mouse: Cre-driver line | (Schwenk *et al.*, 1995) | N/A |
| Mouse: TBA2.1 | Prof. H-U Demuth, Probiodrug, now Vivoryon Therapeutics N.V. (Alexandru *et al.*, 2011) | N/A |
| **Recombinant DNA** | | |
| pSGNluc | (Kuri *et al.*, 2017) | Addgene, Cat.: #186705 |
| LAMP1-GFP | M. Sperveslage | ZMNH, Hamburg |
| RFP-Rab7 | M. Sperveslage | ZMNH, Hamburg |
| **Software and algorithms** | | |
| (Fiji is just) ImageJ | (Schindelin *et al.*, 2012) | http://fiji.sc/; RRID: SCR_002285 |
| Prism Version 9 | GaphPad | https://www.graphpad.com/scientific-software/prism/ |
| **Other** | | |
| Aclar film | Ted Pella, Inc. | Cat.: #10501-10 |
